# Mycobacteria fecal shedding in wild boars (*Sus scrofa*), South-eastern France

**DOI:** 10.1101/670166

**Authors:** Mustapha Fellag, Michel Drancourt, Jean-Lou Marié, Bernard Davoust

## Abstract

The recent recrudescence of tuberculosis in cattle has implicated wild boar as a reservoir and vector of this disease, which led to the investigation of fecal shedding of the causative *Mycobacterium bovis*. In the Provence region of France, wild boars are very abundant, and the current study was carried out to assess the presence of tuberculous and nontuberculous mycobacteria in feces of wild boar population. W e developed an original protocol allowing the fast isolation of mycobacteria by combining a 1%-chlorhexidine decontamination of fecal matter with culture on MOD9 medium. Colonies were identified by matrix-assisted laser desorption/ionization time-of-flight mass spectrometry, combined with DNA sequencing. This protocol yielded no tuberculous mycobacteria among ninety-nine wild boar fecal samples collected in the Provence region. However, non-tuberculous mycobacteria were isolated from five samples (5.05%), including *Mycobacterium peregrinum, Mycobacterium vaccae* and *Mycobacterium setense*, the last species being previously unreported in the wild boar; in addition to two positive samples for *Nocardia* spp. In conclusion, wild boars in southeastern France are not shedding tuberculosis agents, but they could constitute a reservoir of human non-tuberculous mycobacteriosis in selected populations directly exposed to wild boars.

## INTRODUCTION

Bovine tuberculosis is a zoonosis due to *Mycobacterium bovis* (*M*. *bovis*) affecting both livestock and wild mammals ^1^. Bovine tuberculosis is a contagious infection of significant health and economic importance and eradication programs have been implemented in a number of developed countries ^1,2^. These programs yielded a remarkable reduction of the incidence of the disease ^1,3^.

However, the wide distribution of *M. bovis* in wildlife is a major obstacle to the eradication of tuberculosis in cattle ^2,3^. In numerous countries, bovine tuberculosis is maintained by multi-host systems involving cattle and wild mammals ^2–4^. While the role of the red deer (*Cervus elaphus*) remains controversial ^3^, the European badger (*Meles meles*) in Great Britain and Ireland and the wild boar (*Sus scrofa*) in the Iberian Peninsula are acknowledged reservoirs for *M. bovis* ^3,5^. The later species is also recognized as a host for other *Mycobacterium tuberculosis* complex, including *Mycobacterium microti* ^6^ and *Mycobacterium caprae* ^7^. In addition, non-tuberculous mycobacteria that are widely distributed in the environment are not uncommon in wild boars, where they were isolated from lymph nodes ^8^ and feces ^9,10^. These non-tuberculous mycobacteria have been identified as emerging human pathogens, especially in immunocompromised patients ^11,12^.

Accordingly, naturally infected wild boars excrete *M. bovis* mycobacteria by oro-nasal and digestive routes, with inocula above 10^3^ colony-forming units (CFU)/g ^13^. Furthermore, it has been shown that *M. bovis* can persist in soil for several months^14^. Interactions between *M. bovis* and soil-inhabiting amoebae may increase the lifespan of mycobacteria, thus promoting their transmission throughout the environment^15^. Humans and animals may then be contaminated not only by direct contact with infected wild animals, as is the case for hunters handling carcasses^16^, but also from a contaminated environment ^9^.

For epidemiological investigations and the diagnosis of tuberculosis in wild animals, many techniques have been developed, including immunological, serological and molecular biology techniques but culture remains the gold standard technique for the diagnosis of tuberculosis mycobacteria infections ^17^. The lymph nodes are the most frequently used sample for the research of mycobacteria in animals ^17–19^ but this invasive sampling made by well trained staff, is almost limited to dead animals. Therefore, the analysis of feces is an emerging method in wild animals ^13,20–22^ on the model of what has been reported for the routine diagnosis in human patients ^23^.

In France, the first cases of tuberculosis in wild animals were detected in 2001 in the Brotonne forest, Normandy ^2^. Then, monitoring programs for tuberculosis in wild animals were applied and *M. bovis* was detected in badgers, wild boars, red deer, roe deer and red fox in proximity areas of bovine tuberculosis outbreak^4,19,22,24^. One case of tuberculosis was detected in a wild boar in Loir-et-Cher county which had remained free from bovine tuberculosis for more than twenty years^25^. In Southeastern France, there is a lack of information regarding the presence of tuberculosis and non-tuberculous mycobacteria in wild boars, except for information from the national surveillance system for tuberculosis in free-ranging wildlife (Sylvatub) which indicates that wild animals collect in this area are negative ^4^ whereas one study carried-out by serological methods that indirectly detected cases of *M*. *bovis*^26^.

Here, we investigated the presence and the shedding of *Mycobacterium tuberculosis complex* and nontuberculous mycobacteria in feces of wild boars living in Provence region, France by using an original decontamination and culture protocol.

## Results

### Artificially-infected feces samples

Using NA-OH decontamination and culture on Coletsos medium, PBS negative controls remained sterile whereas culture of non-inoculated feces samples were invaded by contaminants. As for feces artificially inoculated with *M. bovis* BCG for ten days, one culture was positive with seven colonies of *M. bovis* BCG confirmed by MALDI-TOF-MS in addition to contaminants, the other two cultures were invaded by contaminants without presence colonies of *M. bovis* BCG. The contaminants were identified by MALDI-TOF-MS as various *Bacillus* species. Using chlorhexidine 1% decontamination and culture on MOD9 medium, PBS negative controls and non-inoculated feces remained sterile. As for feces artificially inoculated with 10^5^ CFUs *M. bovis* BCG, non-frozen feces cultures yielded the first colonies of *M. bovis* BCG after seven and nine days of incubation and more than 100 colonies were observed after 20 days of incubation in one plate and 37 colonies in the second culture plate. Only one *Sphingomonas paucimobilis* contaminant was observed on one of the two cultures. Feces artificially contaminated and frozen at −20°C yielded *M. bovis* BCG colonies after nine-day incubation and after twenty-day incubation and 12 colonies were counted in one plate and only four on the second culture plate, in the absence of any contaminant. These data indicate a sensitivity of 10^3^ CFUs (fresh feces) and 10^4^ CFUs (frozen feces) on MOD9 medium. This method of culture confirmed that feces artificially contaminated with BCG were negative for mycobacteria.

### Culture of wild boar feces

A total of 99 wild boar feces were decontaminated using 1% chlorhexidine and cultured on MOD9 medium. Colonies were observed after 4 to 8 days of incubation. Cultures obtained after one-month incubation were classified into four types: plates containing pure cultures of mycobacteria (n=1); plates containing mycobacteria mixed with contaminants (n=4); plates containing only contaminants (n= 15) including two positive *Nocardia* spp. cultures; and negative-culture plates (n= 79). No further colony was observed after three additional months of incubation (for a total of four-month incubation). None of these 99 samples yielded *M. bovis*, but non-tuberculous mycobacteria were cultivated in five samples which corresponds to a rate of 5.05%. *Mycobacterium vaccae* was isolated in one feces sample collected at military camp A, along with *Mycobacterium peregrinum*in in three samples and *Mycobacterium setense* in one sample, collected at military camp B. These mycobacteria were all identified by MALDI-TOF-MS and confirmed by partial *rpoB* gene sequencing (Table 1). In addition, *Nocardia rhamnosiphila* was identified by 16S rRNA gene sequencing in one sample collected at military camp A and one *Nocardia* spp. isolate representative of a new *Nocardia* species in one sample collected in camp B^27^. MALDI-TOF-MS identified contaminants as *Pseudomonas putida, Pandoraeasputorum, Ralstonia pickettii, Sphingomonas paucimobilis* and *Sphingobacterium faecium*a long with a non-identified fungus.

**Table 1.** Non-tuberculous mycobacteria isolated form wild-boar feces and identified by *rpoB* sequencing; along with *Nocardia* species identified by 16S rRNA sequencing.

| Bacterial species | Number positive | Match of GenBank (%) | GenBank accession | MALDI-TOF score |
| --- | --- | --- | --- | --- |
| *Mycobacterium peregrinum* | 3 | 99 | AY147166.1 | Identified with a high score (2.033) |
| *Mycobacterium setense* | 1 | 100 | HM807426.1 | Identified with a middle score (1.88) |
| *Mycobacterium vaccae* | 1 | 100 | CP011491.1 | Identified with a high score (1.915) |
| *Nocardia rhamnosiphila* | 1 | 100 | KC417288.1 | Not identified |
| Nocardia Strain S-137 | 1 | 98 | CP022088.2 | Not identified |

## DISCUSSION

Culture-based investigation of wild boar feces collected in two geographic areas in the Provence region of France failed to find living tuberculous mycobacteria including *M. bovis*. This observation was made after an original protocol of feces decontamination and culture had been validated using the closely related *M. bovis* BCG strain, in the presence of negative controls which remained negative.

Culture remains the reference method for the routine confirmation of mycobacterial infection in animals^1,28^. The culture protocol herein described allows to detect an inoculum of 10^3^-10^4^ CFUs *M. bovis* which is in the range of that extrapolated from PCR-based observations in wild boars in Spain ^15^. However, contamination is a major limitation of culturing feces. Here, we applied to animal feces a 1% chlorhexidine decontamination and MOD9 medium culture protocol suitable for the recovery of *Mycobacterium tuberculosis complex* in patients on diagnosed with pulmonary tuberculosis ^29,30^. Because the level of contamination of swine feces is very high, an efficient decontamination protocol is essential to isolate and culture mycobacteria from this type of sample^20^. In a first step, we validated this protocol using artificially inoculated feces. Then, we applied it for the first time on animal specimens combined with MOD9 medium protocol and found that this protocol was indeed suitable for the recovery of living mycobacteria from wild boar feces. These results contrasted with the standard NALC-NaOH decontamination and culture on Coletsos medium, which yielded contaminants invading the culture tubes.

Moreover, we observed fewer colonies of *M. bovis* BCG when culturing frozen feces compared to culturing fresh feces. The same observation was previously reported regarding the decreased viability of *M. bovis* after freezing^13^, whereas no effect of freezing at −20°C was reported when culturing *M. tuberculosis* from sputum specimens collected in patients^31^. As for feces, the recommendation is to cultivate fresh feces collected in cattle within less than 96 hours for the isolation and culture of *M. avium* ^21^.

The 1% chlorhexidine decontamination and MOD9 culture protocol enabled the isolation of three non-tuberculous mycobacteria from five different wild boars with a prevalence of 5.05%, in the range of the previously reported 0% to 11.1% prevalence of non-tuberculous mycobacteria in wild boar faeces^10^. In the lymph nodes, prevalence is 16.8% in Spain^8^ whereas no mycobacteria could be detected in lymph nodes in another study in Poland ^32^ the prevalence of mycobacteria in wild boar varies among wild boar population, density and environment. *M. vaccae* was initially cultivated from cows in Austria ^33^ and has previously been isolated in wild boars in Australia ^9^. *M. vaccae* is often considered a non-pathogenic mycobacterium ^34^ and has even been proposed as a vaccine for the treatment of multiresistant tuberculosis^35^. However, cases of pulmonary and skin infections have been described in human diseases^12,33^. Furthermore, *rpoB* gene sequencing confirmed the result obtained by MALDI-TOF and unambiguously ^36^ distinguished *M. peregrinum* from the closely related *M. septicum*. Indeed, the species are not differentiated by 16S rRNA gene sequencing^36^. In animals, *M. peregrinum* has been described as an etiological agent of mycobacteriosis in farmed fish ^37^ was recently isolated in wild boar from Slovenia^18^. It is an opportunistic pathogen previously implicated in surgical site infections and catheter-related infections ^12,38^. Regarding *M. setense* isolated here in one wild boar from military camp B, it has been described for the first time as being responsible for osteitis ^39,40^. Furthermore, three cases have been reported in Italy ^12^ and Iran ^41^. We report here on the first isolation of *M. setense* in wild boars. *M. peregrinum* and *M. setense* are two members of the *Mycobacterium fortuitum* complex, which is including rapidly growing non-tuberculous mycobacteria ^40^ distributed in the aquatic environment ^18,42^. This may suggest the presence of these mycobacteria in the aquatic environments frequented by the wild boar at military camp B.

Moreover, the original culture protocol reported here allowed isolating two species of *Nocardia*. The first isolate was definitely identified as *N. rhamnosiphila* by 16S rRNA gene sequencing. This species, initially identified from a compost mound sample in South Africa ^43^ has never been reported in animals and not in the humans. The second isolate here referred as *Nocardia* strain S-137 is a representative of a hitherto undescribed species: this isolate exhibited a non-identifying MALDI-TOF-MS spectrum, along with a 16S rRNA gene sequence exhibiting only 98% sequence similarity with *N. brasiliensis* strain FDAARGOS. Draft genome sequence and *in silico* DNA-DNA hybridization confirmed strain *Nocardia* S-137 as representative of a new species “*Nocardia suismassiliense*” ^27^.

While the general population is not routinely exposed to wild boars, some selected populations chiefly hunted, are exposed in such way that wild boars do constitute the reservoir of zoonoses, as illustrated by the demonstration of cross-transmission of hepatitis E virus from wild boars to patients in our region ^44,45^.

In conclusion, we found no evidence for living *Mycobacterium tuberculosis* complex, including *M. bovis*, in the feces of wild boars collected during 2015/2016 hunting season in the Provence region of France whereas other actinomycetes including non-tuberculous mycobacteria, were easily cultured. The decontamination-culture protocol reported here can be used for routine, culture-based detection of mycobacteria and related bacteria in wild boar feces samples, a source of previously undescribed microorganisms.

## Materials and methods

### Sample collection

In order to study naturally excreted mycobacteria in wild boar feces, a total of 99 fecal samples were collected during the 2015/2016 hunting season in the Provence region (South-East of France). A total of 79 samples were collected in one military camp, A (Canjuers, Provence, France) (43°38’49’’North; 6° 27’56”East) and 20 samples in another military camp, B (Carpiagne, Provence, France) (43°15’02.1’’North; 5°30’23.6’’East). All samples, collected directly from the animal rectum by hunters were stored at −20 °C until cultured at the latest eight months after collection. No animals were killed specifically for this study and this work did not require animal experiment ethics approval.

### Artificially contaminated feces specimens

*M. bovis* strain BCG was grown at 37°C on Middlebrook 7H10 agar (Becton Dickinson, Pont de Claix, France) supplemented with oleic-albumin-dextrose-catalase (OADC) (Becton Dickinson). The colonies were suspended in a phosphate buffered saline (PBS), then rigorously vortexed using 3-mm sterile glass beads (Sigma-Aldrich, Saint-Quentin-Fallavier, France) and passed five times through a 25-G needle to disperse clustered cells. The homogenized suspensions were then calibrated at 10^7^ CFU/mL by using optical density at 580 nm (Cell Density Meter; Fisher Scientific, Illkirch, France) and calibration curve. Feces of wild boar were inoculated with the suspension of *M. bovis* BCG to a final concentration 10^6^ CFU/g. The artificially contaminated samples were separated into two batches, the first left at room temperature for four hours and the second being frozen at −20°C for one month in order to verify the viability of mycobacteria in frozen feces. The artificially infected samples were then decontaminated using two decontamination methods and cultured using two different culture media, in order to determine the more efficient culture method.

### Decontamination with NALC-NaOH and culture in Coletsos medium

Approximately 1 g of experimentally contaminated feces was placed in a 50-mL Falcon tube complemented with PBS up to 5 mL. An equal volume of N-acetyl-L-cysteine-sodium hydroxide (NALC-NaOH) was added and the tube was vortexed and incubated for 15 min at room temperature with continuous agitation. The remaining volume was completed to 50 mL with a neutralization solution based on a phosphate buffer and centrifuged at 3,000 × g for 20 min. The supernatant was decanted, 1 mL of sterile PBS was added and 250 μL of the pellet-PBS suspension was inoculated, in triplicate, in Coletsos medium (bioMérieux, Craponne, France). PBS and non-inoculated feces were used in parallel as controls for culture to survey for cross-contamination and all tubes were incubated at 37°C.

### Decontamination with chlorhexidine 1% and culture in MOD9 medium

The MOD9 culture medium was prepared as previously described ^29,30^. Briefly, MOD 9 culture medium incorporates for a total volume of 1.000 mL, 5 g of lecithin, 19 g of Middlebrook 7H10 powder, one gram of yeast extract, two grams of glucose, one gram of pancreatic digest of casein, 5 mL of glycerol and 2 mL of Tween 80. All compounds were dissolved in 669 mL of distilled water and the mixture was autoclaved at 121°C for 20 min. Then 150 mL of lamb serum, 100 mL OADC (oleic acid albumin, dextrose, catalase), 20 mL of growth supplement (Becton Dickinson), 50 mL of antibiotic mixture solution (5 mg of vancomycin, 6,000 units of polymyxin B, 600 mg of amphotericin B, 2,400 mg of nalidixic acid, 600 mg of trimethoprim and 600 mg of azlocillin), 4 mL of red food colouring and 0.1g of ascorbic acid were added after the medium was cooled to 55°C. This culture medium has been specially designed for the culture-based recovery of mycobacteria including *Mycobacterium tuberculosis complex*. Decontamination with 1% chlorhexidine was performed according to ^46^ modified by adding filtration as described below. About 1 g of experimentally contaminated feces was deposited into a 50 mL-tube containing 3-mm sterile glass beads, and the volume was completed to 5 mL with sterile PBS. The tube was vortexed and the suspension was filtered with 40 µm cell strainer to remove fibers and other large fecal particles. Two mL of this filtrate were mixed with a triple volume of 1% chlorhexidine, vortexed and incubated for 15 min at room temperature with continuous agitation. The remaining volume was completed to 50 mL with PBS and centrifuged at 3000 X g for 20 min. The supernatant was then decanted, 1 mL of sterile PBS was added, and 100 μL of the pellet-PBS suspension were inoculated, in duplicate, in MOD9 medium. PBS and non-inoculated feces were used in parallel as a negative control for culture to survey for cross contamination, and all plates were incubated at 37°C. This method was further used on to decontaminate and inoculate 99 wild boar feces samples, including 79 samples collected at military camp A, and 20 samples collected at military camp B.

### Artificially contaminated feces culture

The cultures were observed regularly and *M. bovis* strain BCG colonies were counted and identified with the matrix-assisted laser desorption/ionization time-of-flight mass spectrometry (MALDI-TOF-MS) on a Microflex spectrometer (Bruker Daltonics, Bremen, Germany) ^47,48^. Briefly, one loop-full of colonies was removed from the culture and directly applied to the plate and mixed with matrix. For each colony, three different spots were deposited on the mass spectrometer plate and the manipulation was repeated after subculture. The protein profiles were visualized using FlexControl 3.3 software (Bruker Daltonik) and analyzed by the program FlexAnalysis 3.3 (Bruker Daltonik GmbH, Bremen, Germany).

### Wild boar feces culture

After four months incubation at 37°C, colonies observed were subcultured onto the same culture medium and incubated at the same temperature. An initial identification was carried out with MALDI-TOF-MS as previously described ^47,48^. Colonies which remained not identified by MALDI-TOF-MS, were stained by the Ziehl-Neelsen method and examined by using light microscopy. Ziehl-Neelsen-positive colonies were subsequently identified by sequencing PCR-amplified DNA. Genomic DNA was extracted using BIOROBOT EZ1 and the Qiagen Genomic DNA Extraction kit (Qiagen, Courtaboeuf, France) as previously described^49^. Sterile PBS was used as a negative control for extraction.

The 16S rRNA gene PCR amplification and sequencing were performed using the fD1/rP2 primer pair as previously described ^50^. The partial *rpo*B gene PCR amplification and sequencing were performed using the Myco F/Myco R primer pair ^51^. Sequencing was carried-out using the Big Dye Terminator version 3.1 kit (Perkin-Elmer) and the resulting products were recorded with the Biosystems ABI Prism 3130xl as described by the manufacturer (Applied Biosystems, Massachusetts, USA). The obtained nucleotide sequences were assembled using Chromas Pro software, version 1.7 (Technelysium Pty Ltd., Tewantin, Australia) and compared to the GenBank database by similarity search using the BLASTN program (http://www.ncbi.nlm.nih.gov/blast/).

## Declarations

### Acknowledgements

The authors thank the hunters of the military hunting companies of Canjuers and Carpiagne for their very efficient assistance in collecting the samples. This study was supported by IHU Méditerranée Infection, Marseille, France and by the French Government under the «Investissements d’avenir» (Investments for the Future) program managed by the Agence Nationale de la Recherche (ANR, fr: National Agency for Research), (reference: Méditerranée Infection 10-IAHU-03). This work was supported by Région Provence Alpes Côte d’Azur and European funding FEDER PRIMI. M. Fellag benefits a PhD grant from Fondation Méditerranée Infection, Marseille, France. The authors acknowledge Olga Cusack for her expert assistance in editing the manuscript.

### Ethics approval and consent to participate

No Ethics or use committee approval was required since the samples of faeces were collected at a slaughterhouse, on wild boars already killed during hunting according to procedures officially approved by the French authorities (rifle shooting with large caliber bullets) and by authorized hunters. The hunt was not organized for the purpose of sample collection; no animal was killed in the perspective of the present study.

### Consent to publish

Not applicable.

### Availability of data and materials

All data generated or analysed during this study are included in this article.

